# Nutrient starvation activates ECM remodeling gene enhancers associated with inflammatory bowel disease risk in fibroblasts

**DOI:** 10.1101/2024.09.06.611754

**Authors:** Stefano Secchia, Vera Beilinson, Xiaoting Chen, Zi F. Yang, Joseph A. Wayman, Jasbir Dhaliwal, Ingrid Jurickova, Elizabeth Angerman, Lee A. Denson, Emily R. Miraldi, Matthew T. Weirauch, Kohta Ikegami

**Affiliations:** Department of Human Genetics, The University of Chicago, Chicago, Illinois, USA; Department of Biology, Lund University, Lund, 22362, Sweden; Department of Pediatrics, The University of Chicago, Chicago, Illinois, USA; Division of Allergy and Immunology, CCHMC Center for Autoimmune Genomics and Etiology, Cincinnati Children’s Hospital Medical Center, Cincinnati, Ohio, USA; Division of Human Genetics, Cincinnati Children’s Hospital Medical Center, Cincinnati, Ohio, USA; Division of Immunology, Cincinnati Children’s Hospital Medical Center, Cincinnati, Ohio, USA; Division of Gastroenterology, Hepatology, and Nutrition, Cincinnati Children’s Hospital Medical Center, Cincinnati, Ohio, USA; Division of Biomedical Informatics, Cincinnati Children’s Hospital Medical Center, Cincinnati, Ohio, USA; Division of Molecular Cardiovascular Biology, Cincinnati Children’s Hospital Medical Center, Cincinnati, Ohio, USA; Department of Pediatrics, University of Cincinnati College of Medicine, Cincinnati, Ohio, USA

**Author notes:** Institute of Human Biology, Basel, Switzerland (Stefano Secchia). California Institute of Technology, Pasadena, California, USA. **Corresponding author:** Kohta Ikegami, Ph.D.

## Abstract

Nutrient deprivation induces a reversible cell cycle arrest state termed quiescence, which often accompanies transcriptional silencing and chromatin compaction. Paradoxically, nutrient deprivation is associated with activated fibroblast states in pathological microenvironments in which fibroblasts drive extracellular matrix (ECM) remodeling to alter tissue environments. The relationship between nutrient deprivation and fibroblast activation remains unclear. Here, we report that serum deprivation extensively activates transcription of ECM remodeling genes in cultured fibroblasts, despite the induction of quiescence. Starvation-induced transcriptional activation accompanied large-scale histone acetylation of putative distal enhancers, but not promoters. The starvation-activated putative enhancers were enriched for non-coding genetic risk variants associated with inflammatory bowel disease (IBD), suggesting that the starvation-activated gene regulatory network may contribute to fibroblast activation in IBD. Indeed, the starvation-activated gene *PLAU*, encoding uPA serine protease for plasminogen and ECM, was upregulated in inflammatory fibroblasts in the intestines of IBD patients. Furthermore, the starvation-activated putative enhancer at *PLAU*, which harbors an IBD risk variant, gained chromatin accessibility in IBD patient fibroblasts. This study implicates nutrient deprivation in transcriptional activation of ECM remodeling genes in fibroblasts and suggests nutrient deprivation as a potential mechanism for pathological fibroblast activation in IBD.

**HIGHLIGHTS:**

- Serum starvation transcriptionally activates ECM remodeling genes in fibroblasts.
- Fibroblast starvation activates putative distal enhancers associated with ECM remodeling genes.
- Starvation-activated putative enhancers are enriched for inflammatory bowel disease (IBD) risk variants.
- *PLAU* enhancer and expression are activated in IBD intestinal fibroblasts, as in starved fibroblasts.

## INTRODUCTION

Fibroblasts are heterogenous stromal cell types that play critical roles in wound healing, immune response, and fibrosis in vertebrate organ systems ^1,2^. Diverse stimuli activate fibroblasts; these include tissue damage, stress, and immune activation. Activated fibroblasts modulate the extracellular matrix (ECM) by secreting ECM structural proteins and modifying enzymes, thereby altering tissue integrity and intercellular communication. Fibroblast overactivation causes excessive fibrosis contributing to pathology in diseases. Therefore, identifying mechanisms behind fibroblast activation has the potential to contribute to the development of therapeutic approaches to mitigate pathological fibrosis.

Depletion of cell nutrients induces cellular quiescence, a reversible cell cycle arrest state, across various organisms ^3–7^. Nutrient deprivation often results from competition for nutrients among organisms or cells. Interestingly, nutrient deprivation has been observed in pathological tissue microenvironments, such as tumor microenvironments and immune microenvironments ^8–10^. In these contexts, tumor cells or phagocytic immune cells overconsume available nutrients, leading to localized deprivation of nutrients ^8,11^. Fibroblasts are among the major cell types constituting tumor and immune microenvironments ^12,13^. Despite nutrient depletion, however, fibroblasts are often activated within these microenvironments ^12–14^. Activated fibroblasts in pathological microenvironments can contribute to tumor growth, metastasis, chemoresistance, and vascularization through ECM remodeling, immune cell recruitment, cytokine secretion, and metabolic modulation ^12–14^. Whether nutrient deprivation acts as a cue for fibroblast activation remains unclear.

Fibroblast response to nutrient deprivation has been studied *in vitro* by withdrawing serum from the culture medium, a procedure often called "serum starvation". Serum starvation induces cellular quiescence ^7,15^. However, serum-starved fibroblasts remain metabolically active ^16^. Previously studies found that serum starvation alters gene expression programs ^7,17–19^. Specifically, serum starvation results in decreased expression of genes related to cell proliferation and RNA metabolism and increased expression of genes related to tumor suppression, sterol synthesis, immune response, redox, and ECM metabolism ^18^. However, the mechanisms underlying gene expression changes in quiescent fibroblasts remain unclear. Post-transcriptional mechanisms have been proposed to regulate gene expression in quiescent fibroblasts induced by serum starvation or contact inhibition ^19–21^. In contrast, whether quiescent fibroblasts retain robust transcriptional activity remains unclear. Several studies reported that fibroblast quiescence accompanies an increase in histone H4 lysine 20 trimethylation (H4K20me3), a histone modification associated with constitutive heterochromatin, and global chromatin compaction ^22,23^, similar to quiescent starved yeasts ^26,28^. The impact of starvation on transcriptional activity and chromatin states at transcriptional regulatory regions in fibroblasts remains undefined.

Here, we investigate transcriptional and chromatin states in fibroblasts undergoing serum starvation. We report extensive transcriptional activation of ECM remodeling genes in serum-starved fibroblasts, accompanied by the activation of distal putative enhancers. We observe that starvation-activated putative enhancers are enriched for noncoding genetic variants associated with increased inflammatory bowel disease (IBD) risk. We identify *PLAU*, encoding urokinase-type plasminogen activator (uPA), as a starvation-activated, IBD-risk-associated gene. We demonstrate *PLAU* enhancer activation and transcriptional upregulation in inflammatory fibroblasts in IBD patient intestines. Together, our data nominate local nutrient deprivation as a candidate mechanism for fibroblast activation contributing to IBD.

## RESULTS

### Large-scale transcriptional upregulation during fibroblast starvation

To define the transcriptional state of fibroblasts undergoing nutrient deprivation, primary dermal fibroblasts (GM07492) maintained in 10% serum (“proliferation”) were subjected to 1% serum for 48 hours (“starvation”) (**Fig. 1A**). Starved cells were then subjected to a recovery phase with 10% serum for 24 hours ("recovery"). The S/G2 cell-cycle population was 72% decreased at the end of the starvation phase relative to the proliferation phase and then almost completely restored (96%) at the end of the recovery phase (**Fig. S1**), consistent with the previous observation that serum depletion induces cellular quiescence ^7^. To determine genes transcribed during proliferation, starvation, and recovery, we performed Bru-seq, in which nascent transcripts are labeled with 5-bromouridine (BrU) in cell culture, purified by anti-BrU immunoprecipitation, and quantified by high-throughput sequencing ^30^. We labeled transcripts for 2 hours, either during proliferation, at the end of starvation, or at the end of recovery (**Fig. 1A**). Bru-seq signals covered both exons and introns, as expected for nascent transcripts (**Fig. 1B**), and were highly concordant between replicates (**Fig. S2A, B**). We identified extensive differences of Bru-seq signals among the three growth conditions. Proliferating and starved fibroblasts differed in transcriptional states at 1,002 genes (394 up- and 608 down-regulated in starvation relative to proliferation), while starved and recovering fibroblasts differed in transcription at 2,090 genes (1,126 up- and 964 down-regulated in recovery relative to starvation) (**Fig. 1C, D; Fig. S2C**). In total, we identified 2,336 unique genes with differential transcription between proliferation and starvation, starvation and recovery, or both, representing 11.7% of the 19,955 protein-coding genes analyzed (**Fig. 1E**). Thus, transcriptional states differed extensively between proliferating, starved, and recovering fibroblasts.

**Figure 1.**
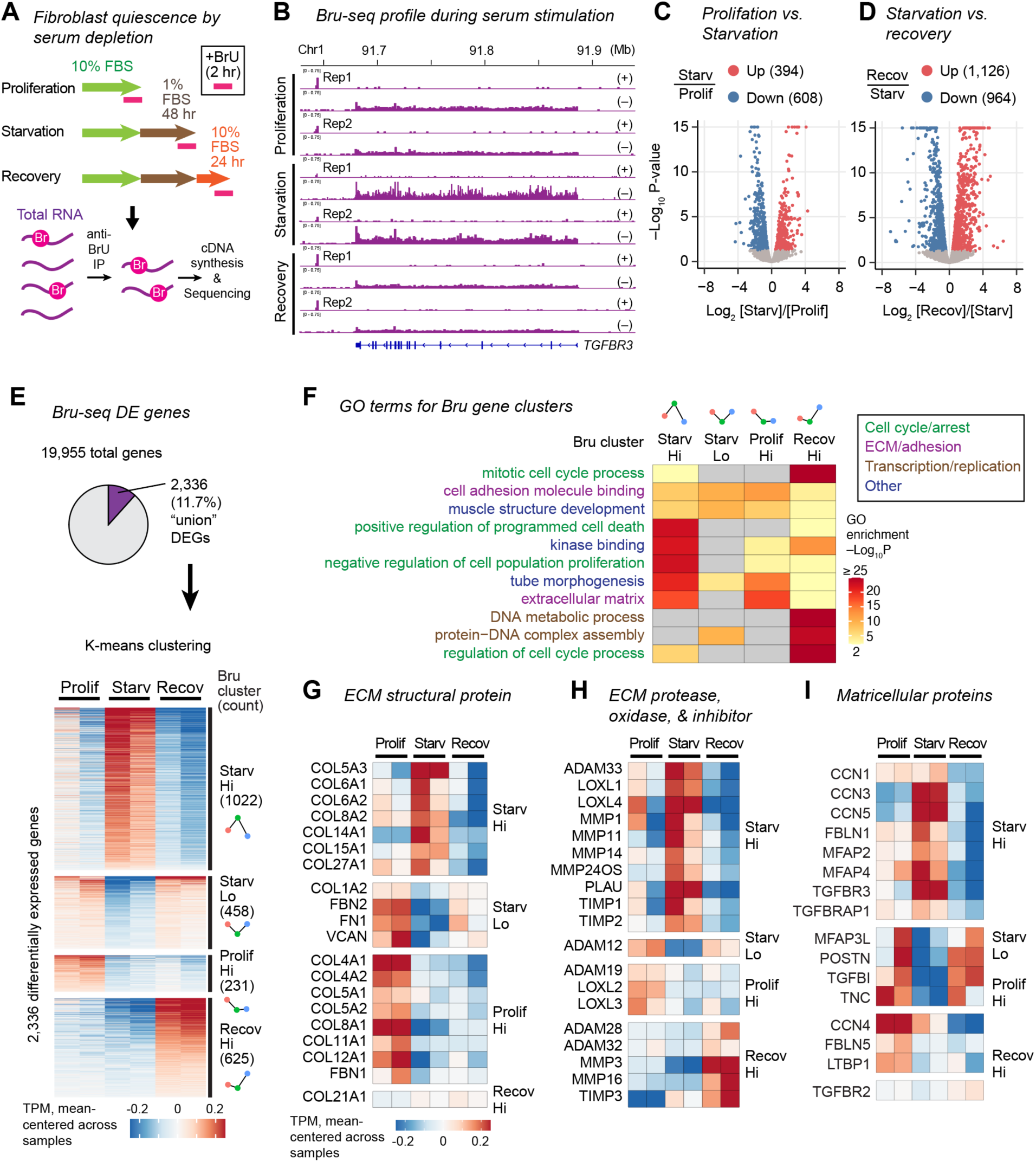
Serum starvation upregulates transcription of ECM remodeling program in fibroblasts. **A.** Bru-seq nascent transcript profiling in fibroblasts undergoing proliferation, starvation, and recovery from starvation. **B.** Bru-seq signal tracks at *TGFBR3* gene upregulated during starvation. **C.** Volcano plot comparing Bru-seq transcript abundance between proliferating and starved fibroblasts. Parenthesis, gene count. **D.** Same as C but showing difference between starved and recovering fibroblasts. **E.** Clustering of differentially transcribed genes. Top, the fraction of union differential transcribed genes over total genes. Bottom, normalized transcript level of differentially transcribed genes grouped by the dynamics of gene expression. Small plots along the right Y-axis represent mean expression values within cluster and serve as icons in this and subsequent figure panels. **F.** GO terms enriched in differentially transcribed Bru-seq gene clusters. **G–I.** Normalized transcript level of differentially transcribed genes.

To identify patterns of transcriptional changes, we clustered the 2,336 differentially expressed genes based on the transcriptional dynamics. This analysis identified four distinct patterns of transcriptional dynamics (**Fig. 1E; Table S1**). The Starv-Hi gene cluster (1,022 genes) consisted of genes upregulated during starvation and downregulated during recovery. The Starv-Hi cluster was the largest cluster, representing 44% of all differentially expressed genes. The remaining three clusters, Prolif-Hi (231 genes), Recov-Hi (625 genes), and Starv-Lo (458 genes), had the lowest expression during starvation. Their peak transcription occurred during proliferation (Prolif-Hi), recovery (Recov-Hi), or both proliferation and recovery (Starv-Lo). Thus, Bru-seq revealed large-scale transcriptional activation during fibroblast starvation, contrary to the prevailing hypothesis for cellular quiescence.

### Upregulation of the ECM-remodeling gene program during starvation

To identify biological pathways specifically activated during starvation, we investigated Gene Ontology (GO) terms overrepresented within each of the four Bru-seq gene clusters (**Fig. 1F**). Starv-Hi genes were strongly enriched for the “extracellular matrix” (ECM) GO term. In addition, Starv-Hi genes were overrepresented for the “negative regulation of cell population proliferation” GO term, consistent with the reduction of cell cycle activity. Interestingly, the ECM GO term was also enriched among Prolif-Hi genes. Because Bru-seq gene clusters were mutually exclusive, this result suggested that distinct ECM-related genes were upregulated in proliferating fibroblasts or starved fibroblasts.

Upregulation of ECM-related genes in starved fibroblasts was interesting because enhanced ECM gene expression is a typical feature of activated, but not quiescent, fibroblasts. We thus investigated specific ECM-related genes that were upregulated or downregulated in starved fibroblasts. We observed a complex alteration of collagen gene expression during starvation (**Fig. 1G**). For example, we observed a switch in transcribed fibril-forming collagen genes (upregulated: *COL5A3* and *COL27A1*; downregulated: *COL1A2*, *COL5A1*, *COL5A2*, and *COL11A1*), upregulation of filament collagen genes (*COL6A1, COL6A2*), and downregulation of network-forming collagen genes (*COL4A1*, *COL4A2*, *COL8A1*). Additionally, major non-collagen ECM components, fibronectin (*FN1*) and fibrillin (*FBN2*), were downregulated during starvation. We further observed alteration of ECM modifying enzyme expression during starvation (**Fig. 1H**). The genes upregulated during starvation included ECM-degrading proteinases (*MMP1*, *MMP11*, *MMP14, PLAU*), ECM proteinase inhibitors (*TIMP1*, *TIMP2*), and ECM-crosslinkers (*LOXL1* and *LOXL4*). Moreover, matricellular protein genes were differentially expressed during starvation (**Fig. 1I**). The upregulated matricellular protein genes included *CCN3*, *CCN5*, and *TGFBR3*, while downregulated matricellular protein genes included *POSTN*, *TGFBI*, and *LTBP1*. The downregulation of the myofibroblast marker *POSTN* ^27^ suggested that starved fibroblasts were distinct from myofibroblasts. Together, the extensive transcriptional alteration of genes encoding ECM components, ECM modifying enzymes, and matricellular proteins suggested that serum starvation upregulated a gene expression program promoting ECM remodeling in fibroblasts.

### Pervasive gains of H3K27ac at distal putative enhancers during starvation

To define the mechanism underlying the transcriptional activation during starvation, we performed ChIP-seq for histone H3 lysine 27 acetylation (H3K27ac), a chromatin modification associated with active regulatory elements, in proliferating, starved, and recovering fibroblasts matching the Bru-seq experimental conditions. (**Fig. 2A, B**). The H3K27ac profiles were highly concordant between biological replicates but differed extensively among the three growth conditions (**Fig. S3A, B**). We identified 7,846 differentially H3K27-acetylated genomic sites between proliferation and starvation (3,353 gained and 4,493 lost in starvation relative to proliferation) (**Fig. 2C**), and 9,747 differentially H3K27-acetylated sites between starvation and recovery (5,796 gained and 3,951 lost in recovery relative to starvation) (**Fig. 2D**). In total, 14,392 unique genomic sites were differentially H3K27-acetylated among these three conditions, representing 24% of the total 60,704 H3K27ac sites (**Fig. 2E**).

**Figure 2.**
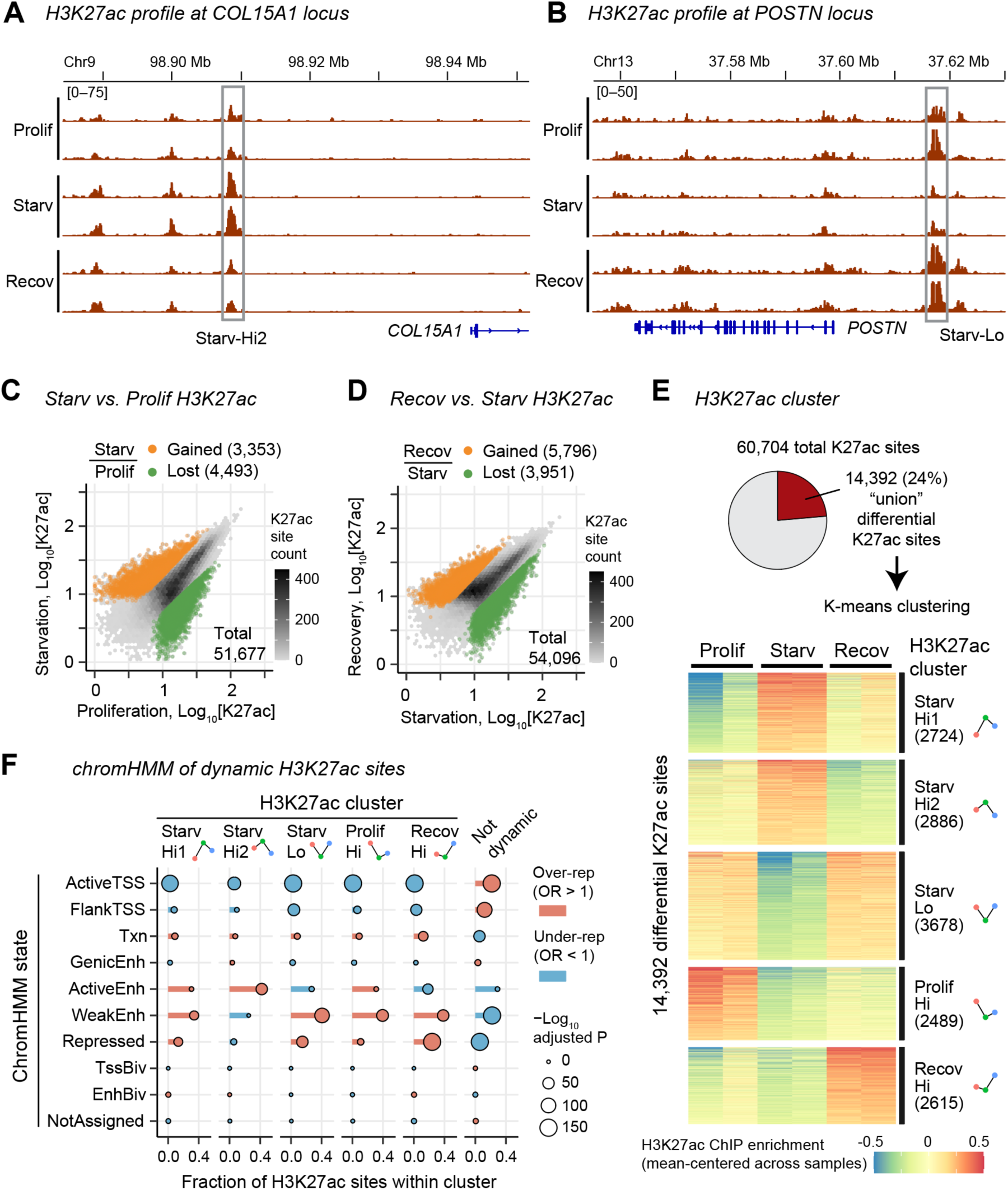
Extensive gains of H3K27ac at distal putative enhancers in starved fibroblasts. **A, B.** Representative H3K27ac ChIP-seq signal tracks at *COL15A1* (**A**) and *POSTN* (**B**) loci. Differentially H3K27 acetylated regions are indicated. **C, D.** Scatter plot comparing normalized ChIP-seq read coverage between proliferation and starvation (**C**) and between starvation and recovery (**D**) at H3K27ac peak locations identified. Parenthesis indicates H3K27ac site count. **E.** Clustering of union differentially H3K27ac-marked sites. Top: Proportion of union differentially H3K27ac- marked sites among all H3K27ac sites. Bottom: K-means clustering of the differentially H3K27ac-marked sites. Small plots along the right Y-axis represent mean H3K27ac scores within cluster and serve as icons in this and subsequent figure panels. **F.** Enrichment of chromatin states within differentially H3K27ac-marked clusters.

Clustering of the 14,392 differential H3K27ac sites identified five dynamic patterns of H3K27ac changes, similar to the transcription dynamics identified in Bru-seq (**Fig. 2E; Table S2**). The Starv-Hi1 (2,724 sites) and Starv-Hi2 (2,886 sites) H3K27ac clusters showed peak H3K27ac levels during starvation (e.g. *COL15A1* locus, **Fig. 2A**). These two clusters differed in that Starv-Hi1 had higher H3K27ac levels during recovery, while Starv-Hi2 had higher H3K27ac levels during proliferation. The remaining four clusters had the lowest H3K27ac levels during starvation and peaked during proliferation (Prolif-Hi, 2,489 sites), recovery (Recov-Hi, 2,615 sites), or both proliferation and recovery (Starv-Lo, 3,678 sites; e.g. *POSTN* locus, **Fig. 2B**). Starv-Hi1 and Starv-Hi2 H3K27ac sites accounted for 39% of all differential H3K27ac sites, indicating extensive H3K27ac gains during starvation.

To define the genomic characteristics of the dynamic H3K27ac sites, we investigated the chromatin state of these genomic locations, using the chromatin state annotation in proliferating skin fibroblasts ^33^. The dynamic H3K27ac clusters were enriched for “active enhancers” (Starv-Hi2) or “weak enhancers” (Starv-Hi1, Starv-Lo, Prolif-Hi, and Recov-Hi) and depleted in transcription start sites (TSSs) (**Fig. 2F**). In contrast, non-dynamic H3K27ac sites were enriched for active TSSs and depleted in active or weak enhancer sites. These data suggested that the extensive alteration of H3K27ac primarily occurred at putative distal enhancers, not TSSs, during starvation.

### The gain of H3K27ac accompanies transcriptional upregulation of ECM-related genes during starvation

We tested whether changes in H3K27ac were associated with changes in transcription during starvation. We linked each H3K27ac site to a gene if the site was located within the gene body or within 100 kb centered around the TSS. Starv-Hi1 and Star-Hi2 H3K27ac sites were strongly overrepresented at Starv-Hi genes (**Fig. 3A**). Conversely, H3K27ac sites with low H3K27ac levels during starvation (Starv-Lo, Prolif-Hi, and Recov-Hi H3K27ac sites) were enriched for genes with low transcription during starvation (Starv-Lo, Prolif-Hi, and Recov-Hi genes). Interestingly, Starv-Hi1 and Star-Hi2 H3K27ac sites were also overrepresented, although modestly, at Prolif-Hi genes.

**Figure 3.**
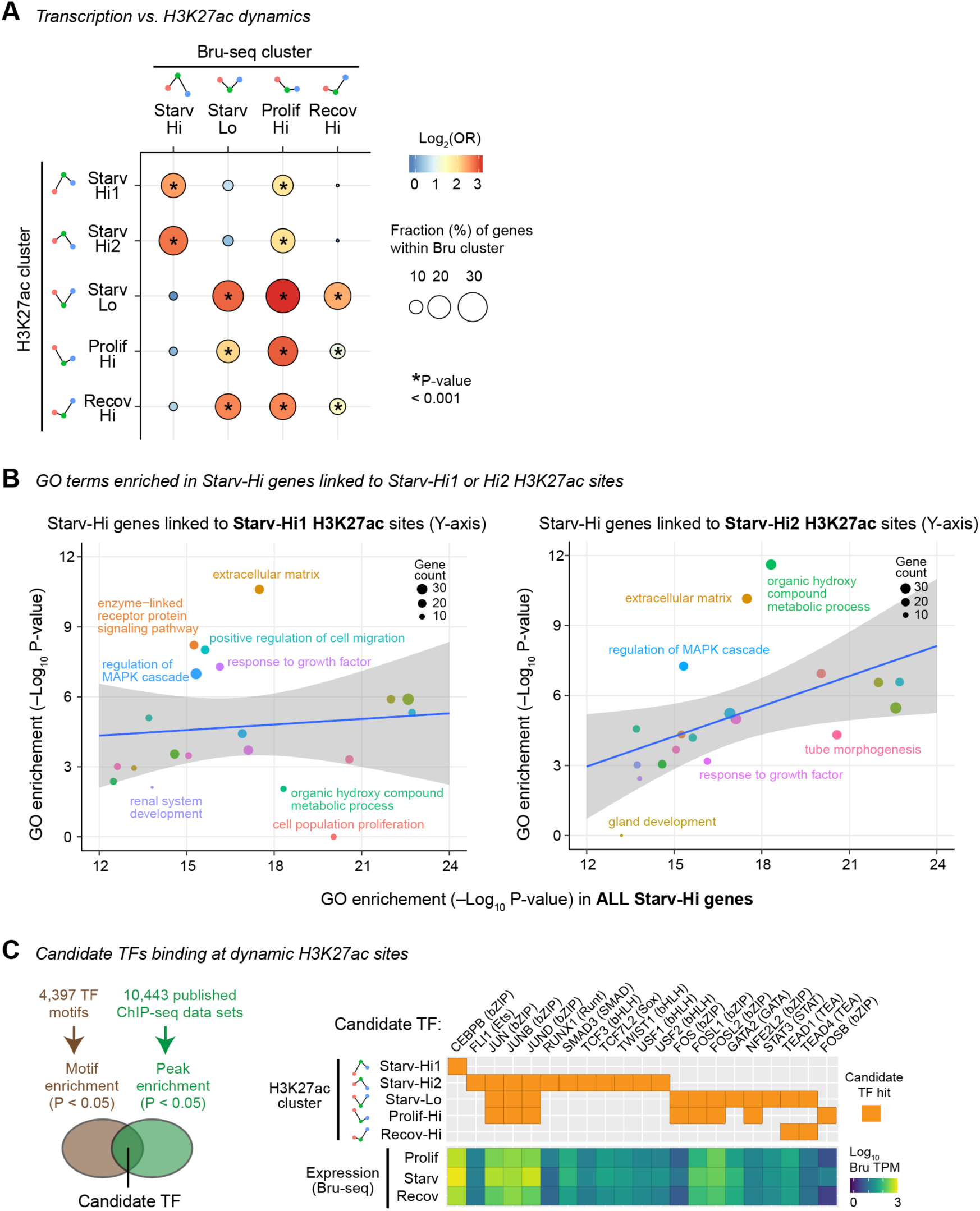
Gains of H3K27ac occur at upregulated ECM-related genes in starved fibroblasts. **A.** Fraction of Bru-seq cluster genes with indicated H3K27ac cluster sites. **B.** GO terms enriched in Starv-Hi genes linked to Starv-Hi1 H3K27ac sites (left) or Starv-Hi2 H3K27ac sites (right) versus GO terms enriched in all Starv-Hi genes. Each circle represents each GO term, with the circle size indicating the number of Starv-Hi genes linked to Starv-Hi1 or Hi2 H3K27ac sites. Line, linear least squares. Shade, 0.95 confidence interval. GO terms above or below the 95% interval range of the linear model (text labeled) suggest over- or under-representation, respectively, in the H3K27ac-linked subset of Starv-Hi genes than predicted. **C.** Candidate TFs binding to H3K27ac dynamic cluster sites. Left: Candidate TFs are those whose DNA motifs are enriched within DNase hypersensitive sites in the dynamic H3K27ac sites and whose public ChIP-seq peaks (in any cell types) are enriched in the dynamic H3K27ac sites. Right: Candidate TFs identified for each dynamic H3K27ac site cluster (top) and their expression levels measure by Bru-seq (bottom).

To identify gene pathways potentially upregulated by the starvation-dependent increase of H3K27ac at Starv-Hi1 or Starv-Hi2 sites, we investigated GO terms overrepresented in Starv-Hi genes associated with Starv-Hi1 or Starv-Hi2 H3K27ac sites. The subset of Starv-Hi genes linked to Starv-Hi1 or Starv-Hi2 H3K27ac sites were most strongly overrepresented for the “extracellular matrix” GO term among all GO terms associated with Starv-Hi genes (**Fig. 3B**). In contrast, the cell proliferation and cell death-related GO terms (e.g. “positive regulation of programmed cell death” and “negative regulation of cell population proliferation”) were not overrepresented within the subsets of Starv-Hi genes linked to Starv-Hi1 or Starv-Hi2 H3K27ac sites. Thus, Starv-Hi1 and Starv-Hi2 H3K27ac sites might act as enhancers for ECM-related gene transcription during starvation.

### Candidate transcription factors operating at starvation-activated putative enhancers

We next identified candidate transcription factors (TFs) operating at the dynamic H3K27ac sites. We identified TFs whose binding sites were overrepresented within dynamic H3K27ac clusters by surveying 10,443 published ChIP-seq datasets using the RELI program ^29^. The identified candidate TFs were further narrowed down to TFs whose binding motifs were also enriched in local accessible chromatin regions within or immediately adjacent to the H3K27ac sites (**Fig. 3C**). The single resulting candidate TF overrepresented at Starv-Hi1 H3K27ac sites was CEBPB, an important transcriptional activator for inflammatory responses ^30^. The candidate TFs overrepresented at Starv-Hi2 H3K27ac sites were bHLH TFs (TWIST1, TCF3, USF1, USF2), JUN TFs (JUN, JUNB, JUND), TCF7L2, FLI1, RUNX1, and SMAD3. Among those, TWIST1 has been implicated in cancer-associated fibroblast activation ^27,29,31,34,36^, while SMAD3 is a critical regulator of myofibroblast transformation downstream of TGF-beta ^38,40^. Both Starv-Lo and Prolif-Hi H3K27ac sites were enriched for AP-1 TF binding sites (JUN, JUND, JUNB, FOSL1, FOSL2), which are dominant pro-proliferation TFs ^42^. Finally, the candidate TFs for Recov-Hi H3K27ac sites were TEAD1 and TEAD4, which are oncogenic transcription factors for cell proliferation ^43,44^. These analyses suggested that distinct sets of TFs regulate putative enhancers during proliferation, starvation, and recovery, with potential important roles for CEBPB, bHLH TFs, JUN TFs, TCF7L2, FLI1, RUNX1, and SMAD3 in regulating starvation-activated putative enhancers.

### Promoter-associated H3K4me3 remains largely unaltered in starved fibroblasts

The investigation of H3K27ac dynamics suggested that the chromatin states of promoters were largely unchanged during starvation. To further examine the promoter chromatin states, we performed ChIP-seq for promoter-associated histone H3K4 trimethylation (H3K4me3) in proliferating, starved, and recovering fibroblasts. We confirmed high consistency of H3K4me3 enrichment between biological replicates (**Fig. S3D, E**). We found that only 99 sites (0.6%) gained and 74 sites (0.4%) lost H3K4me3 levels during starvation compared with proliferation, among the total 16,507 H3K4me3-marked sites found in these two conditions (**Fig. 4A**). Similarly, only 50 sites (0.3%) gained and 129 sites (0.8%) lost H3K4me3 levels during recovery compared with starvation (**Fig. 4B**). In total, only 320 sites of the total 17,095 H3K4me3 sites (1.9%) were differentially H3K4-trimethylated among the three conditions (**Fig. 4C**). Thus, unlike H3K27ac sites, a small subset of H3K4me3 sites were affected by serum starvation in fibroblasts.

**Figure 4.**
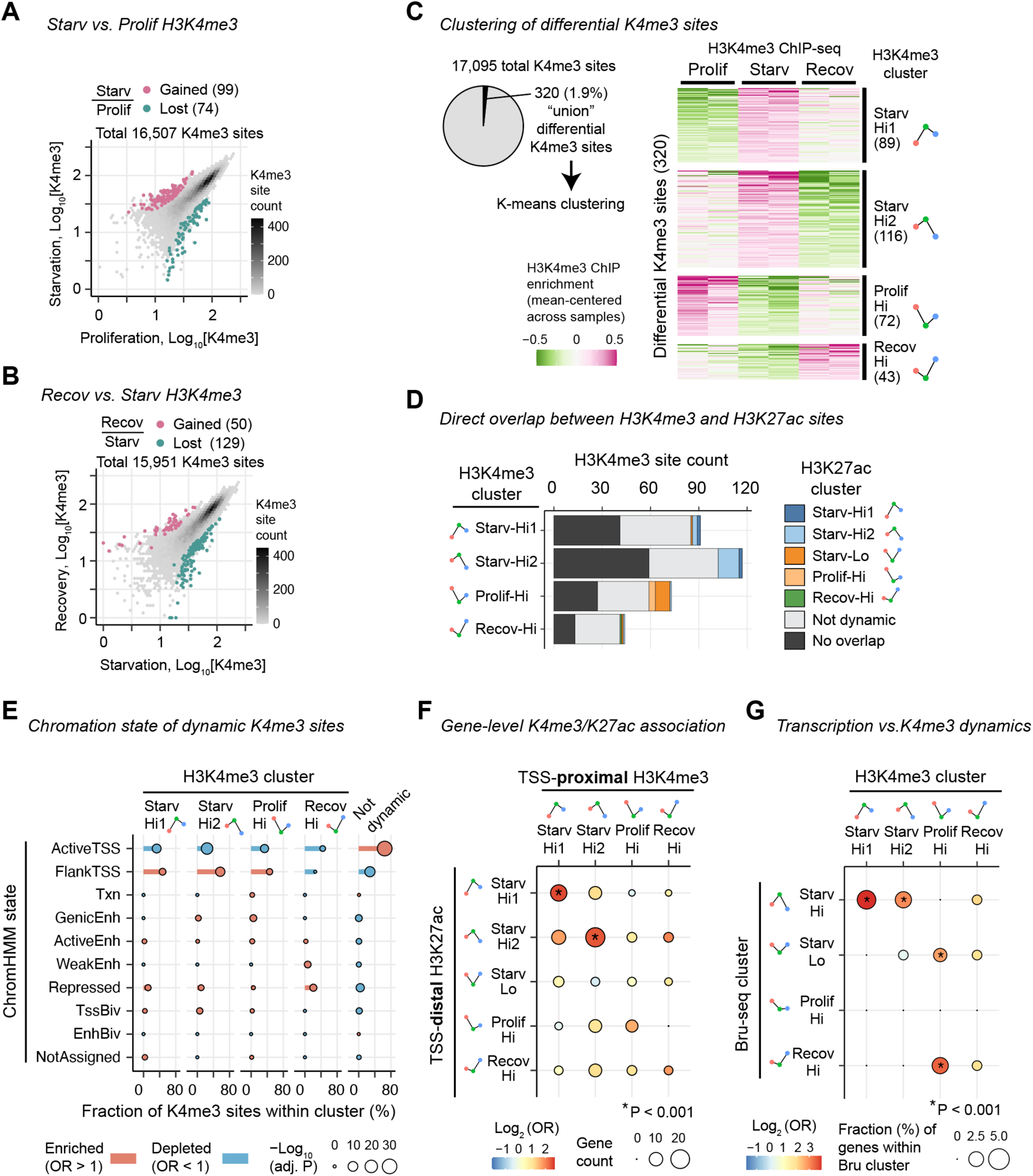
Starvation caused limited alteration to promoter-proximal H3K4me3 states. **A, B.** Scatter plot comparing normalized ChIP-seq read coverage between proliferation and starvation (**A**) and between starvation and recovery (**B**) at H3K4me3 peak locations identified. Parenthesis indicates H3K4me3 site count. **C.** Clustering of union differentially H3K4me3-marked sites. Left: Proportion of union differentially H3K4me3-marked sites among all H3K4me3 sites. Right: K-means clustering of the differentially H3K4me3-marked sites. Small plots along the right Y-axis represent mean H3K4me3 scores within cluster and serve as icons in this and subsequent figure panels. **D.** Number of dynamic H3K4me3 sites, stratified by the state of direct overlap with H3K27ac sites. **E.** Enrichment of chromatin states within differentially H3K4me3-marked site clusters. **F.** Gene-level co-occurrence between TSS-proximal H3K4me3 sites and TSS-distal H3K27ac sites. **G.** Fraction of Bru-seq cluster genes linked to indicated H3K4me3 cluster sites.

We investigated the dynamics of the 320 differential H3K4me3 sites by clustering. These sites were grouped into four clusters, displaying similar patterns to the differential H3K27ac sites (**Fig. 4C; Table S3**). The two largest clusters, Starv-Hi1 (89 sites) and Starv-Hi2 (116 sites), showed the highest H3K4me3 levels during starvation, analogous to Starv-Hi1 and Starv-Hi2 H3K27ac sites. Starv-Hi1 and Hi2 H3K4me3 sites differed in that Starv-Hi1 had higher H3K4me3 levels during recovery, while Starv-Hi2 had higher levels during proliferation. The Prolif-Hi (72 sites) and Recov-Hi (43 sites) clusters exhibited high H3K4me3 levels during proliferation or recovery, respectively, similar to the Prolif-Hi and Recov-Hi H3K27ac sites. Dynamic H3K4me3 sites did not include a group analogous to the Starv-Lo H3K27ac sites, which had high H3K27ac levels in both proliferation and recovery phases. Given the similarity in H3K4me3 and H3K27ac dynamics, we asked whether the dynamic sites of these two modifications overlapped each other. We observed only 2 of the 89 Starv-Hi1 H3K4me3 sites (2%) overlapped Starv-Hi1 H3K27ac sites, while 13 of the 116 Starv-Hi2 H3K4me3 sites (11%) overlapped Starv-Hi2 H3K27ac sites (**Fig. 4D**). Thus, the changes of H3K4me3 occurred largely at genomic locations distinct from the dynamic H3K27ac sites.

We examined the genomic features of the dynamic H3K4me3 sites by comparing them with chromatin state annotations derived from proliferative fibroblasts (**Fig. 4E**). Starv-Hi1, Starv-Hi2, and Prolif-Hi H3K4me3 sites were overrepresented in regions annotated as "TSS flanking" regions but underrepresented in "Active TSS" regions. The "TSS flanking" annotation represents regions enriched within +/– 1 kb of active TSS but outside of the TSS proper (+/– 200 bp) ^33^. Recov-Hi H3K4me3 sites were not strongly overrepresented in any of the chromatin states. In contrast, non-dynamic H3K4me3 sites were strongly overrepresented at active TSSs. These findings suggested that the limited starvation-associated changes in H3K4me3 occurred primarily at regions proximal to TSSs.

We hypothesized that the changes of H3K4me3 at TSS-proximal sites might be associated with the changes of H3K27ac at TSS-distal sites at the same gene loci. To test this hypothesis, we linked H3K4me3 sites to genes using the same algorithm employed for H3K27ac sites and investigated genes linked to TSS-proximal H3K4me3 sites (overlapping Active or Flanking TSS chromatin states) and genes linked TSS-distal H3K27ac sites (not overlapping above TSS states). Indeed, we observed that the genes linked to Starv-Hi1 TSS-proximal H3K4me3 sites had Starv-Hi1 TSS-distal H3K27ac sites more frequently than expected by chance (**Fig. 4F**). Similarly, we observed a significant association between Starv-Hi2 proximal H3K4me3 sites and Starv-Hi2 distal H3K27ac sites. Thus, the H3K4me3 gains at select TSS-proximal sites were associated with distal H3K27ac gains during starvation.

We next asked whether the dynamic H3K4me3 sites were associated with differentially transcribed genes identified by Bru-seq. We found that both Starv-Hi1 and Starv-Hi2 H3K4me3 sites were significantly enriched at Starv-Hi genes (4% and 3% of Starv-Hi genes respectively; **Fig. 4G**). However, the frequencies of these associations were much lower than the frequencies of the association between Starv-Hi1 and Starv-Hi2 H3K27ac sites and Starv-Hi genes (22% and 29% of Starv-Hi genes respectively; **Fig. 3A**). These results indicated that the limited TSS-proximal H3K4me3 gains did not explain a substantial fraction of transcription upregulation during starvation and suggested that distal H3K27ac gains had a larger influence on this transcription upregulation without accompanying proximal H3K4me3 gains.

### IBD-associated genetic variants are overrepresented within starvation-gained H3K27ac sites

Our analysis so far revealed that the gains of H3K27ac were associated with transcriptional activation of ECM-related genes in starved fibroblasts. Given the role of ECM remodeling in pathological fibrosis, we tested the hypothesis that starvation-gained H3K27ac sites are enriched for non-coding genetic variants associated with diseases. To test this hypothesis, we investigated whether the dynamic H3K27ac sites were overrepresented for disease-associated genetic variants identified by genome-wide association studies (GWAS), using the RELI algorithm ^29^. Strikingly, genetic variants associated with inflammatory bowel disease (IBD), including Crohn’s disease and ulcerative colitis, were specifically overrepresented within Starv-Hi2 H3K27ac sites, but not other dynamic H3K27ac sites including Starv-Hi1 H3K27ac sites (**Fig. 5A**). Starv-Hi2 H3K27ac sites were also overrepresented for variants associated with other conditions, such as male pattern baldness, heel bone mineral density, and systolic blood pressure, and monocyte percentage of white blood cells. Only weak enrichment of disease-associated variants, if any, was found in other dynamic H3K27ac site clusters.

**Figure 5.**
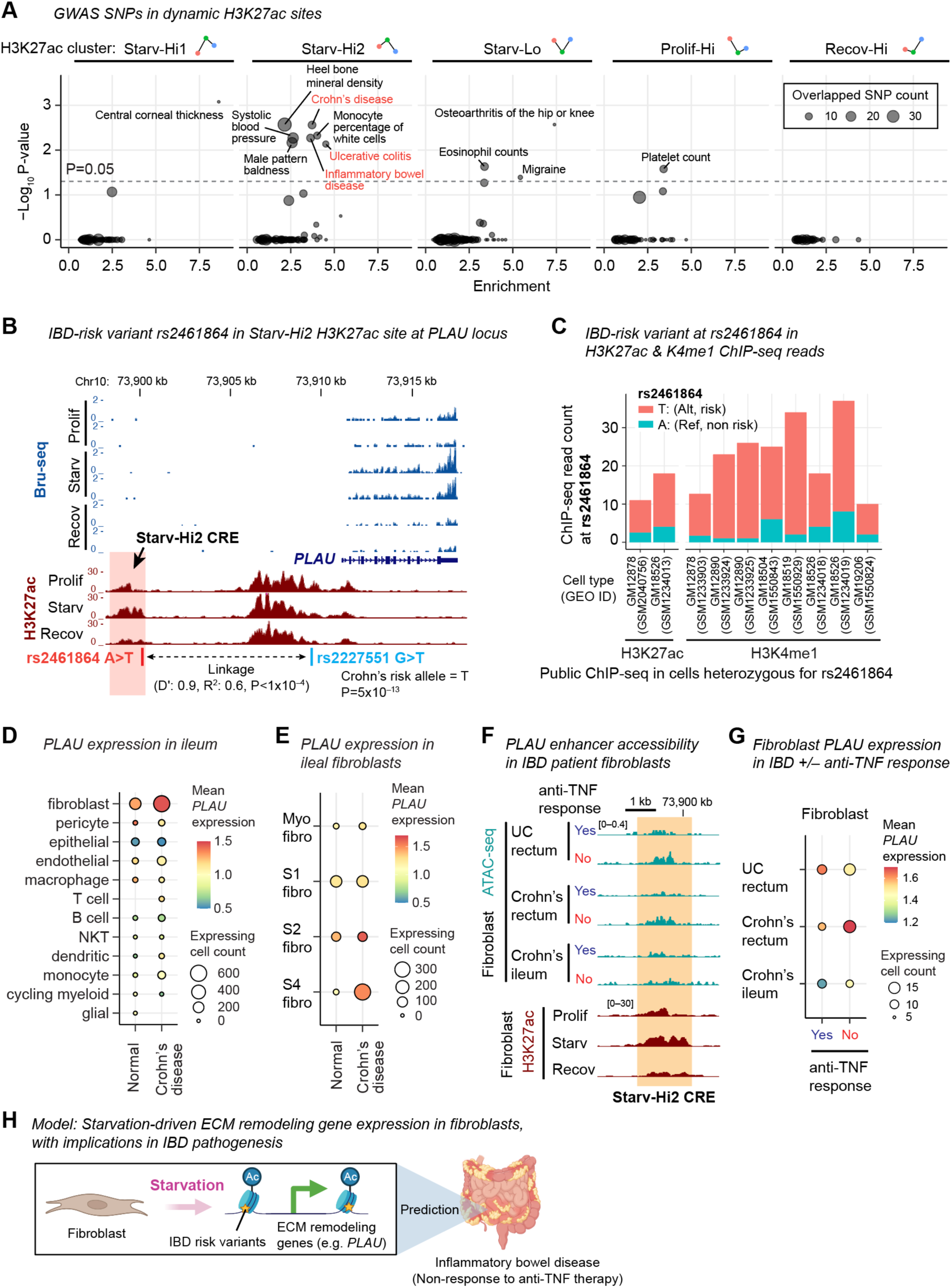
Starvation-activated H3K27ac sites are enriched for IBD-risk variants. **A.** GWAS SNP enrichment in H3K27ac clusters. **B.** Locations of IBD-risk SNP rs2461864 within Starv-Hi2 H3K27ac CRE at the *PLAU* locus. NES, normalized effect size for eQTL. eQTL P, eQTL P-value. D’, normalized linkage strength (0 to 1). R^2^, correlation coefficient of determination. LD P, chi-square P-value for linkage. **C.** ChIP read count with T or A variant at rs2461864 in public H3K27ac and H3K4me1 ChIP-seq datasets heterozygous for the allele. **D.** *PLAU* expression level and *PLAU*-expressing cell count by cell types in normal ilea and Crohn’s disease patient ilea. Data are from Elmentaite et al. 2020. **E.** Same as D, but data are stratified by fibroblast cell types. **F.** ATAC-seq signals at Starv-Hi2 CRE in intestinal fibroblasts in IBD patients responsive or non-responsive to anti-TNF therapy (indicated as Yes or No, respectively). The H3K27ac data (bottom) is the same as data in **B** and shown as a reference. Data are from Wayman et al. 2024. See **Figure S5** for signal quantification. **G.** *PLAU* expression level and *PLAU*-expressing cell count in IBD patients responsive or non-responsive to anti-TNF therapy. Data are from Wayman et al. 2024. **H.** Summary. Starvation activates ECM remodeling genes via putative enhancer activation in fibroblasts. Starvation-activated putative enhancers are enriched for IBD risk variants.

### The *PLAU* locus harbors an IBD-associated genetic risk non-coding variant within a starvation-gained H3K27ac site

The overrepresentation of IBD-related variants within Starv-Hi2 H3K27ac sites was interesting because aberrant fibroblast activation is associated with IBD pathogenesis ^35,37^. Therefore, we investigated IBD-related GWAS variants within Starv-Hi2 H3K27ac sites near Starv-Hi genes. We identified that the *PLAU* gene locus harbored the non-coding variant rs2461864 (A>T) within a Starv-Hi2 H3K27ac site, which we named “Starv-Hi2-CRE” (*cis*-regulatory element) for convenience (chr10:73,898,518-73,900,174 in hg38) (**Fig. 5B**; **Table 1**). *PLAU* is a starvation-activated Starv-Hi gene and encodes urokinase-type plasminogen activator uPA, which generates plasmin, a potent protease for fibrinolysis and ECM-remodeling ^39,41^. The Starv-Hi2-CRE variant rs2461864 exhibited strong positive linkage disequilibrium with *PLAU* TSS-proximal SNP rs2227551 (G>T), which was originally identified as a Crohn’s disease risk SNP (P=5×10^-13^ for allele T) ^45,46^. The Starv-Hi2-CRE variant rs2461864 also showed positive linkage disequilibrium with three other SNPs near *PLAU* (rs2227564, rs2688607, rs2688608), all of which were associated with increased IBD risks (P=7×10^-10^; P=6×10^-11^; P=3×10^-10^, respectively) (**Table 1**) ^46–48^. These data indicated that the alternative "T" allele of the Starv-Hi2-CRE variant rs2461864 was the risk-associated allele for IBD. The risk alleles of Starv-Hi2-CRE variant rs2461864 and other IBD risk-associated SNPs at the *PLAU* locus were individually associated with high *PLAU* expression in fibroblasts in eQTL studies (**Table 1**) ^50^. These data suggested the association between the Starv-Hi2-CRE variant rs2461864 risk allele, *PLAU* expression, and IBD risk.

We next tested whether the risk allele of Starv-Hi2-CRE variant rs2461864 was associated with the activity of this putative enhancer. We observed that, in various independent ChIP-seq experiments performed on cells heterozygous for the rs2461864 allele, the risk allele was strongly overrepresented in H3K27ac and H3K4me1 ChIP reads compared to the non-risk allele (**Fig. 5C**). This supported the positive correlation between the risk allele and the Starv-Hi2-CRE activity. We next investigated the genotype of rs2461864 in the GM07492 fibroblast line used in the serum starvation experiment. All ChIP-seq and input sequencing reads spanning this SNP harbored the risk allele (**Fig. S4**), suggesting that GM07492 was homozygous for the risk allele at rs2461864. This was consistent with the high frequency of the risk allele among European descendants (0.73 at rs2461864), from which this cell line was derived. This analysis suggested that starvation-induced signaling acted on the IBD-risk allele at Starv-Hi2 CRE to transcriptionally activate *PLAU* expression in fibroblasts. Together, activation of Starv-Hi2-CRE might contribute to activation of *PLAU* in fibroblasts in IBD in a genotype-dependent manner.

### *PLAU* is upregulated in inflammatory fibroblasts in IBD patients’ intestines

We tested the hypothesis that *PLAU* was upregulated in intestinal fibroblasts in IBD patients, similar to starved fibroblasts in culture. We analyzed published single-cell RNA-seq data from small intestines (ileum) of pediatric Crohn’s disease patients and normal individuals ^52^. We observed both increased *PLAU* expression and increased *PLAU*-expressing cell fraction in fibroblasts, but not other cell types, in Crohn’s disease versus normal ileal biopsies (**Fig. 5D**). Among the Crohn’s disease intestinal fibroblasts, *PLAU* expression was most strongly upregulated within an inflammatory fibroblast population designated as “Stromal 4 (S4)” ^54,56^ (**Fig. 5E**), which is strongly expanded in Crohn’s disease (**Fig. S5A**). Thus, *PLAU* was strongly upregulated in intestinal inflammatory fibroblasts derived from IBD patients, similar to serum-deprived cultured fibroblasts.

Given the association between *PLAU* expression and Starv-Hi2 CRE activity in starved fibroblasts, we predicted that the Starv-Hi2 CRE might be activated in IBD-derived fibroblasts. We examined published single-cell ATAC/RNA-seq multiome data from ileal and rectal biopsies from pediatric IBD patients, responsive (defined by endoscopic healing) or non-responsive (defined by endoscopic ulcers) to standard-of-care anti-TNFα antibody therapy ^58^. We hypothesized that non-responsive IBD patients might exhibit the activated Starv-Hi2-CRE, given the report that *PLAU* expression was increased in inflamed over non-inflamed colorectal tissues ^60^. Indeed, we found increased chromatin accessibility for the Starv-Hi2-CRE in the fibroblasts of non-responsive Crohn’s disease and ulcerative colitis rectal biopsies compared with responsive counterparts (**Fig. 5F; Fig. S5B**). An enhanced accessibility was not observed in non-responder Crohn’s disease ileal biopsies. We next examined whether the increased accessibility of the Starv-Hi2-CRE accompanied transcriptional activation of *PLAU*. We found enhanced expression of *PLAU* in fibroblasts of non-responsive Crohn’s rectal biopsies compared with responsive counterparts, while such enhanced expression was not apparent in ulcerative colitis rectal biopsies potentially due to sparse read coverage (**Fig. 5G**). Collectively, these results supported the hypothesis that the Starv-Hi2-CRE promotes *PLAU* expression within fibroblasts in certain contexts of inflamed IBD.

In conclusion, this study unveiled starvation-driven transcription of ECM remodeling genes in fibroblasts and nominates nutrient starvation as a candidate mechanism for ECM remodeling in fibroblasts in IBD via activating IBD-risk predisposed distal enhancers (**Fig. 5H**).

## DISCUSSION

Our observation that serum-starved fibroblasts are transcriptionally highly active and gain H3K27ac at thousands of putative distal enhancers challenges the paradigm that quiescent cells are transcriptionally dormant ^22,26,28^. Previous studies reported that heterochromatin-associated H4K20me3 increases at retrotransposons and ribosomal RNA genes in serum-starved fibroblasts ^23^. Our investigation suggest that distal enhancers undergo distinct regulatory mechanisms than genomic repetitive regions to orchestrate ECM remodeling gene transcription during starvation.

This study demonstrates that *PLAU* as a starvation-driven gene in fibroblasts upregulated by a starvation-activated putative enhancer (Starv-Hi2-CRE). We further unveil that the Starv-Hi2-CRE possesses an IBD risk-associated noncoding variant (rs2461864) that promotes this putative enhancer activity. *PLAU* encodes the urokinase-type plasminogen activator uPA, which converts plasminogen to plamin, a potent protease for fibrinolysis and ECM degradation, widely implicated in cancer metastasis^39^. Increased *PLAU* and uPA expression in IBD patient intestines has been known for decades ^49,51^. Recent investigation pinpointed that the cellular source of *PLAU* expression in IBD patient intestines is neutrophils ^60^ and fibroblasts ^53,54^, although the upstream signaling has been unknown. Our study nominates that local nutrient deprivation as a candidate mechanism for *PLAU* upregulation in intestinal fibroblasts, which may be further amplified by the IBD risk-associated variant within the Starv-Hi2-CRE putative enhancer (**Fig. 5H**). While a recent study found that *PLAU* genetic deletion ameliorates a mouse model of colitis ^60^, the exact pathogenic role of *PLAU* remains controversial ^55^ and requires further investigation.

Our study reports an increased chromatin accessibility of the Starv-Hi2-CRE in fibroblasts from IBD patients non-responsive to the standard-of-care anti-TNFα antibody treatment. A previous study reported that the anti-TNF therapy does not reduce the stricture complication in IBD and that this complication is associated with upregulated expression of ECM-related genes in intestines ^57^. It is worth investigating whether the potential starvation-driven activation of ECM remodeling genes in IBD fibroblasts contributes to anti-TNF non-response and stricture complications in IBD.

A limitation of this study is that serum starvation, as used in this study, does not model specific pathological nutrient-deprived conditions *in vivo*, which themselves are largely unknown. Our serum starvation condition is likely deprived of certain growth factors, cytokines, enzymes, hormones, lipids, and minerals included in fetal bovine serum, but not amino acids, vitamins, and glucose, which were included in the basal culture medium. The effect of specific nutrient deprivation on fibroblast activity *in vivo* warrants further investigation.

Production of ECM components, matricellular proteins, and ECM-modifying enzymes is a widely recognized feature of activated proliferative fibroblasts. In this regard, our observation that starved quiescent fibroblasts actively transcribe these ECM-related genes is interesting. A similar upregulation of ECM-related genes in starved fibroblasts was reported in the analysis of mature RNAs ^7,18,19^. Our observation that starved fibroblasts and proliferating fibroblasts express distinct ECM-related genes suggests that both entry into and exit from starvation involves ECM remodeling. We speculate that local nutrient starvation in microenvironments might promote ECM remodeling gene transcription, similar to cultured fibroblasts. A recent study reports that nutrient starvation promotes cytokine production in various cells in culture and suggested that starvation may serve as a inflammation initiator ^60^. We speculate that nutrient starvation also serves as an initiator of ECM remodeling, thereby contributing to various disease pathology such as fibrosis, metastasis, and IBD.

## ACKNOWLEDGEMENTS

We thank Sebastian Pott, Ivan Moskowitz, members of the Moskowitz laboratory for critical feedback on this study. We thank the genomics core facility at the University of Chicago for their assistance. This work is supported by NIH grant R21/R33 AG054770 (K.I.), R21HG012423 (K.I.), CCHMC Research Innovation and Pilot grant (K.I.), U24HG013078 (M.T.W.), R01AI148276 (M.T.W.), P30AR070549 (M.T.W.), R01HG010730 (M.T.W.), and CCHRF Academic Research Committee (ARC) Award #53632 (M.T.W.), CCHRF ARC grant #53671 (E.R.M., L.A.D.), and Center for Pediatric Genomics (CpG) grant (E.R.M., L.A.D., K.I.), U01AI150748 (E.R.M., M.T.W.), R01AI153442 (E.R.M.), R01AI173314 (E.R.M., M.T.W.), R01AI148276 (L.C.K., M.T.W.), P30DK078392 (L.A.D.).

## AUTHOR CONTRIBUTIONS

Conceptualization, S.S. and K.I.; methodology, S.S. and K.I.; formal analysis, S.S., V.B., X.C., J.A.W., Z.F.Y. and K.I.; investigation, S.S., V.B., and K.I.; resources, L.A.D. M.W., and E.R.M.; data curation, K.I., S.S., J.A.W., I.J., E.A., M.W., E.R.M., L.A.D.; writing – original draft, S.S., K.I.; writing – review & editing, K.I., M.W., E.R.M., L.A.D; visualization, S.S., V.B., K.I.; supervision, M.W., E.R.M., L.A.D., and K.I.; project administration, M.W., E.R.M., L.A.D., K.I.; funding acquisition, L.A.D., M.T.W., E.R.M., K.I.

## METHODS

### Cell culture

Human skin fibroblast cell line GM07492 (Coriell Cell Repository) was grown in MEM Alpha medium (Life Technologies, catalog number 12561-056) with 10% FBS (Sigma) and 1% Penicillin/Streptomycin mix (Corning) and incubated at 37°C with 5% CO_2_. This was the proliferation growth condition. For serum starvation, the proliferation culture medium was replaced with MEM Alpha with 0.1% FBS, 1% Penicillin/Streptomycin, at 37°C with 5% CO_2_ for 48 hours. For recovery, the 48-hour starved cells were cultured with the proliferation medium at 37°C with 5% CO_2_ for 24 hours.

### Cell cycle analysis

Fibroblast cells were trypsinized, resuspended in 500 µL of PBS + 0.01% Tween 20, and then fixed in 1 mL ethanol at –20°C for at least 2 hours. The cells were washed 3 times with 1 mL FACS buffer (PBS supplemented with 2% heat-inactivated FBS, 1 mM EDTA, 0.01% Tween 20) and stained with DAPI. The cells were then washed twice with FACS buffer and then analyzed for DAPI intentsity in LSR-Fortessa 4-9 HTS bench top flow cytometer (BD Biosciences). DAPI intensity for at least 10,000 individual cells were recorded for each sample.

### Bru-seq

Bru-seq was performed by following the protocol developed elsewhere with modifications ^30^. To label RNAs in GM07492 fibroblast cells in culture, we added 5-Bromouridine (BrU, Sigma #850187, dissolved in PBS at 50 mM) directly to the culture medium at the final concentration of 2 mM and incubated cells at 37°C with 5% CO_2_ for 2 hours. This 2-hour labeling was performed during proliferation, at 46 to 48 hours into starvation, and at 22 to 24 hours into the recovery phase. Two hours later, the culture medium was removed, and cells were harvested with Trizol LS for RNA extraction (Invitrogen #10296010). Purified total RNA was treated with TurboDNase (Ambion AM18907) and fragmented by Fragmentation Buffer (Ambion AM8740). BrU-incorporated RNA fragments were immunoprecipitated with mouse monoclonal anti-BrdU antibody 3D4 (BD Biosciences 555627 Lot 7033666) and used to construct DNA sequencing libraries using NEBNext Ultra II Directional RNA Library Prep kit (New England Biolabs E7760). DNA libraries were sequenced on an Illumina HiSeq 4000 machine for single-end 50-nt sequencing. Bru-seq was performed in two biological replicates per condition. The experiment IDs are KI658 and KI662 for proliferation, KI659 and KI663 for starvation, and KI661 and KI665 for recovery.

### ChIP-seq

ChIP-seq in GM07492 fibroblasts were performed during proliferating, at 48 hours into starvation, and at 24 hours into the recovery phase. The detail ChIP-seq procedures in GM07492 fibroblasts was described in our previous publication ^61^. Cells were crosslinked in 1% formaldehyde for 15 min at room temperature. Cell extract from one million cells was incubated with antibodies in a 200-μL reaction for 12 hours or longer at 4°C. Antibodies used in ChIP are: mouse monoclonal anti-H3K27ac antibody (Wako MABI0309, Lot 14007, RRID:AB_11126964; 2 μL per IP) and mouse monoclonal anti-H3K4me3 antibody (Wako MABI0304, Lot 14004, RRID:AB_11123891; 2 μL per IP). Immunocomplex was captured by Protein G-conjugated magnetic beads and washed. Immunoprecipitated DNA was reverse-crosslinked and used to construct high-throughput sequencing libraries using NEBNext Ultra DNA Library Prep Kit (New England Biolabs, E7370). DNA libraries were processed on an Illumina HiSeq machine for single-end 50-nt sequencing. ChIP-seq was performed in two biological replicates per condition. The experiment IDs for H3K27ac ChIP-seq are KI351 and KI422 for proliferation, KI374 and KI401 for starvation, and KI379 and KI402 for recovery. The experiment IDs for H3K4me3 ChIP-seq are KI353 and KI403 for proliferation, KI376 and KI404 for starvation, and KI381 and KI405 for recovery. The experiment IDs for input sequencing are KI384 for proliferation, KI396 for starvation, and KI397 for starvation.

### Bru-seq analysis

Bru-seq reads were aligned to the human hg38 reference genome with the Gencode v38 basic gene annotation using STAR version 2.7.9 ^62^ with the default alignment parameters except using "-- clip3pAdapterSeq *AGATCGGAAGAGCACACGTCTGAACTCCAGTCA* --sjdbGTFfeatureExon *gene*" parameters. From raw read counts per transcript, TPMs (transcripts per million) were calculated in R as follows: TPM = 10^6^ x RPK/sum(RPK), where RPK = [read count] x 1/[transcript length in kb], and sum(RPK) is the sum of RPKs for all transcripts. Raw read counts were used for differential gene expression analyses between proliferation (KI658 and KI662) and starvation (KI659 and KI663) and between starvation (KI659 and KI663) and recovery (KI661 and KI665) using DESeq2 in R. Protein-coding genes with adjusted *p*-value <0.05 were considered differentially expressed genes. This yielded 2,336 non-overlapping set (i.e. "union") of differentially expressed genes. To cluster genes based on transcription levels, a matrix of TPMs for the union of differentially expressed genes for sample replicates was processed using the *kmeans* function with k=4, which captured major transcription dynamics observed (**Table S1**). High correlation of Bru-seq TPM between replicates, a principal component analysis (PCA) showing the expected sample clustering, and differentially expressed gene counts are shown in **Fig. S2**.

### ChIP-seq analysis

ChIP-seq reads were mapped to the hg38 human reference genome using Bowtie2 ^63^ with the default “--sensitive” parameter. Reads with MAPQ score greater than 20 were used in downstream analyses. We generated per-base coverage of 200 bp-extended aligned reads normalized by sequencing depth, and this was used for visualization of ChIP-seq genomic signal distribution. To identify ChIP enriched "peak" locations, the aligned reads from biological replicates of ChIP and the corresponding input were processed by MACS2 (v2.1.0) ^64^ with "call peak -g hg --nomodel --extsize *200* --SPMR --bdg --call-summits" parameter set. This also generated input-normalized, replicate-combined per-base fold enrichment signal scores, which were used in visualization of replicate-combined genomic signals. Peak locations with the fold-enrichment score greater than 5 were kept for subsequent analyses. For each histone modification type, a union of peak locations from the three culture conditions (proliferation, starvation, and recovery) were generated by merging peak locations if neighboring peak edges were located within 50 bp. Final union peak locations were established after removing peak locations overlapping ENCODE blacklist regions (see **Public Dataset**). For each peak location in the union set, we computed the sum of the per-base coverage of 200 bp-extended ChIP-seq reads for each biological replicate (unnormalized read coverage) (**Table S2, Table S3**). From this dataset, normalized read coverage was computed as follows: normalized base coverage = 10^6^ x RPK/sum(RPK), where RPK = [base count] x 1/50 x 1/[peak size in kp], and sum(RPK) is the sum of RPKs for all peak locations. To identify differentially ChIP-enriched regions between conditions, DESeq2 analysis was performed on unnormalized read coverage within the subset of the union peaks that had peaks in either or both of the two conditions under comparison. Peak locations with the absolute log_2_ fold change > 1 and adjusted *p*-value < 0.05 were identified as differentially enriched. For clustering, the subset of the union peak locations that were identified as differentially enriched between proliferation and starvation, between starvation and recovery, or both were used. In clustering, a matrix of normalized base coverage for sample replicates was processed using the *kmeans* function with k=5 for H3K27ac or k=4 for H3K4me3, which captured major chromatin dynamics observed. High correlation of ChIP-seq coverage at peak regions between replicates, a PCA showing the expected sample clustering, and differentially enriched site counts are shown in **Fig. S3**.

### Chromatin state analysis

We used the 18-state ChromHMM annotation for normal human dermal fibroblast (E126, NHDF) developed by the Roadmap Epigenomics Project (see **Public Dataset**) ^33^. The 18 states were consolidated into the following 9 states for simplicity: 1. Active TSS (Active TSS), 2. TSS Flanking (TSS Flank, TSS Flank Upstream, TSS Flank Downstream), 3. Transcription (Transcription), 4. Genic Enhancer (Genic Enhancer 1, Genic Enhancer 2), 5. Active Enhancer (Enhancer Active 1, Enhancer Active 2), 6. Weak Enhancer (Weak Enhancer), 7. Repressed (ZNF/Repeats, Heterochromatin, Polycomb Repressed, Weak Polycomb Repressed, Quiescent), 8. Bivalent TSS (Bivalent TSS), and 9. Bivalent Enhancer (Bivalent Enhancer). The genomic overlap between ChIP-seq peak locations and chromHMM states was computed using the Bedtools *intersect* function ^65^. In cases when multiple states overlapped a peak location, a single state with the largest overlap was assigned. The over- and under-representation of each chromatin state within each dynamic cluster of H3K27ac or H3K4me3 sites over all sites was computed by Fisher’s exact test with the p-value adjusted for multiple testing using the Benjamini-Hochberg procedure.

### Joint analysis of Bru-seq and ChIP-seq

Each H3K27ac or H3K4me3 ChIP peak was assigned to one gene or none using the following algorithm, in order. If a peak overlapped the gene body of a single gene, this peak was assigned to that gene. If a peak was in the gene body of multiple genes, this peak was assigned to the gene whose TSS was closest to the peak. If a peak was located less than +/– 50 kb of the TSS of a single gene, this peak was assigned to that gene. If a peak was located less than +/– 50 kb of TSSs of multiple genes, this peak was assigned to the gene whose TSS was closest to the peak. Other peaks were not assigned to a gene. The over- and under-representation of dynamic H3K27ac or H3K4me3 sites within Bru-seq cluster genes over all genes was computed by Fisher’s exact test with the p-value adjusted for multiple testing using the Benjamini-Hochberg procedure.

### Gene Ontology (GO) analysis

Metascape ^66^ was used to evaluate the enrichment of GO Biological Processes, GO Molecular Functions, GO Cellular Components, and KEGG pathway terms with the default enrichment parameters. For the comparison of term enrichment between different Bru-seq clusters, gene lists were provided as a multiple gene list file. Similarly, a multiple gene list file containing all Starv-Hi genes and a subset of Starv-Hi genes associated with Starv-Hi1 or Hi2 H3K27ac sites was analyzed.

### Candidate TF identification

DNA motif analysis was performed on DNase I hypersensitive sites (DHS) located within H3K27ac ChIP-seq peak locations. We first obtained a list of 3.5 million DHSs from 438 human cell and tissue types ^67^ (see **Public Dataset**). We then extended H3K27ac ChIP-seq peak locations 50 bp to each side to include potential linker regions adjacent to H3K27ac nucleosomes. We next identified all DHS centers (1 bp) located within the extended H3K27ac sites. Using FIMO ^68^ with the human Cis-BP motif database (version 2.0) ^69^, we identified all DNA motifs located within +/–25 bp of the DHSs within the H3K27ac peak locations. We applied Fisher’s exact test to identify DNA motifs enriched within DHSs of H3K27ac clusters over all H3K27ac sites with the p-value adjusted for multiple testing using the Benjamini-Hochberg procedure. We considered a DNA motif with the adjusted p-value < 0.05 and odds ratio > 1 as overrepresented. Independently, we used RELI ^29^ to assess enrichment of any of 10,443 published ChIP-seq datasets within a cluster of H3K27ac locations over all H3K27ac peak locations, with p-value < 0.05 as a threshold.

### GWAS analysis

Enrichment of GWAS variants within dynamic H3K27ac peak locations was computed using RELI ^29^. Briefly, we extracted phenotypes to analyze by filtering GWAS catalog v1.0.2 entries with five or more independent tag SNPs (r^2^<0.2) under European ancestry for each phenotype. RELI analysis was then performed on the resulting 763 phenotypes using the default settings described in our original paper ^29^. The genome-wide significance *P*-value of select SNPs at the *PLAU* locus was obtained from EBI GWAS catalog as of June 1, 2024 (**Public Dataset**). The eQTL significance *P*-value and normalized effect size (NES) in cultured fibroblasts were obtained from GTEx Analysis Release V8 (dbGaP Accession phs000424.v8.p2) database in GTEx Portal (**Public Dataset**). The degree of correlation between rs2461864 and rs2227551 was computed using LDpair Tool in LDlink ^70^ (**Public Dataset**) using the GRCh38 High Coverage genome build and "All Populations" as the target calculation population. The allelic imbalance of public ChIP-seq reads at rs2461864 was analyzed using MARIO ^29^ using a database of 1,058 publicly available ChIP-seq data in lymphoblastoid cell lines described in our previous paper ^71^.

### Single-cell RNA-seq and ATAC-seq analysis

We obtained the normalized single-cell RNA-seq dataset for terminal ileum of pediatric Crohn’s disease and matched healthy controls ^52^ from the Gut Cell Atlas website (**Public Dataset**). The data were processed using the *SingleCellExperiment* R package. The single-cell ATAC-seq data for inflamed (anti-TNF-alpha refractory) or non-inflamed (anti-TNF-alpha responsive) pediatric Crohn’s disease recta, pediatric ulcerative colitis patient recta, and pediatric Crohn’s disease ilea are previously described ^58^. The normalized pseudo-bulk ATAC-seq read coverage was used to obtain coverage at Starv-Hi2-CRE.

### Data visualization

Genomic signal tracks were visualized using IGV ^72^. All other data were visualized using the *ggplot* package in R.

### Public Dataset and tools

#### ENCODE blacklist regions in hg38

https://www.encodeproject.org/files/ENCFF356LFX/

#### ChromHMM annotation

https://egg2.wustl.edu/roadmap/data/byFileType/chromhmmSegmentations/ChmmModels/core_K27ac/jointModel/final/E126_18_core_K27ac_hg38lift_mnemonics.bed.gz

#### DNaseI hypersensitive sites

https://zenodo.org/record/3838751#.YXIeONnMImo

#### GWAS Catalog

https://www.ebi.ac.uk/gwas/

#### GTEx Portal

https://gtexportal.org/

#### LDpair Tool

https://ldlink.nih.gov/

#### RELI

https://github.com/WeirauchLab/RELI

#### MARIO

https://github.com/WeirauchLab/MARIO

#### Pediatric Crohn’s disease scRNA-seq in Gut Cell Atlas

https://cellgeni.cog.sanger.ac.uk/gutcellatlas/final_pediatric_object_cellxgene_c.h5ad

#### IGV (Integrative Genomics Viewer)

https://igv.org/doc/desktop

---

**Supplementary Figure S1.**
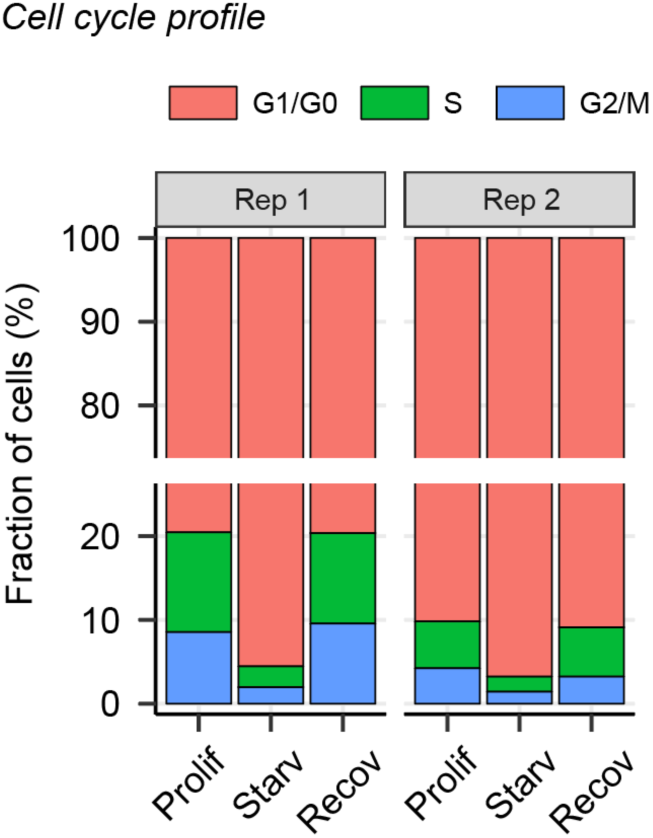
Cell-cycle profile of fibroblasts undergoing proliferation, serum-starvation, and recovery from starvation.

**Supplementary Figure S2.**
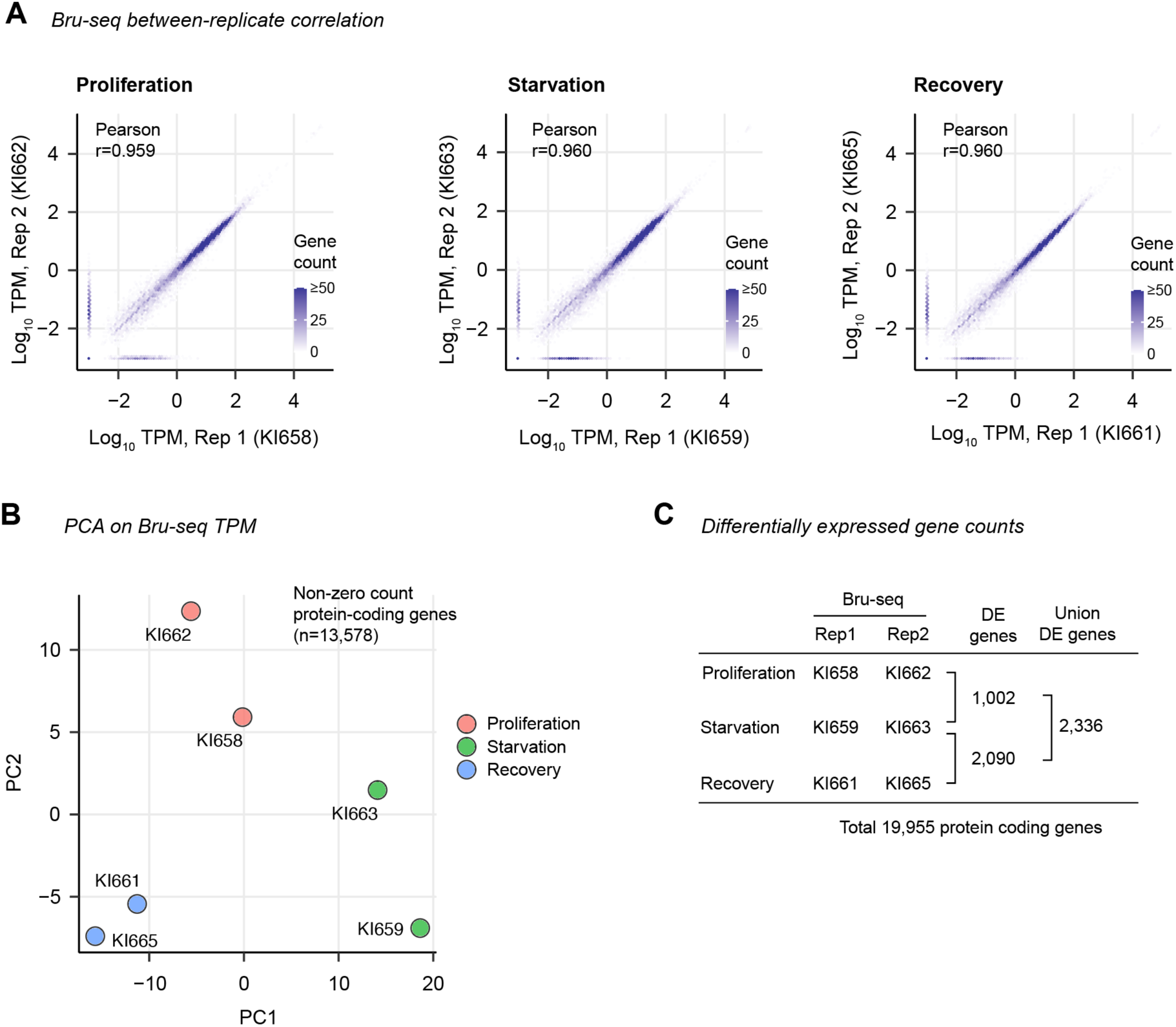
Bru-seq TPM correlation between replicates. **A.** Distribution of Bru-seq TPMs at 19,955 protein-coding genes is plotted as a two-dimensional histogram between biological replicates, with color indicating the number of genes in each bin. **B.** PCA on Bru-seq TPM for 13,578 protein-coding genes that have non-zero TPMs for at least one of the six samples. **C.** Sample IDs for Bru-seq data and the number of differentially expressed genes.

**Supplementary Figure S3.**
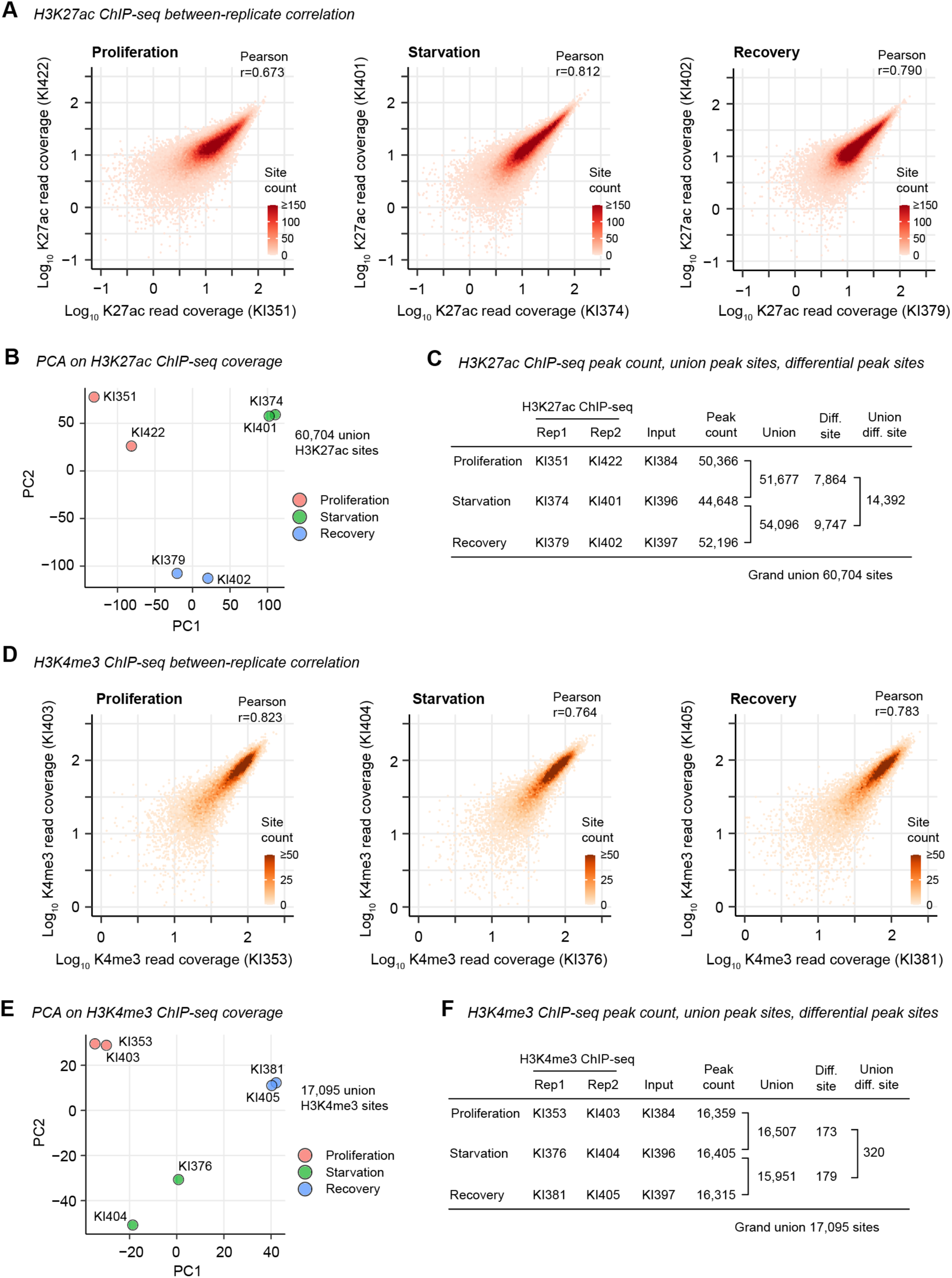
ChIP-seq read quality assessment. **A.** Distribution of H3K27ac ChIP-seq read coverage at the union of H3K27ac ChIP-seq peaks is plotted as a two-dimensional histogram between biological replicates, with color indicating the number of sites in each bin. **B.** PCA on H3K27ac ChIP-seq read coverage at union H3K27ac peak regions. **C.** H3K27ac ChIP-seq and input data IDs used in this study, and H3K27ac peak counts for each condition. **D, E, F.** Same as **A**, **B**, **C**, respectively, but H3K4me3 ChIP-seq data are shown.

**Supplementary Figure S4.**
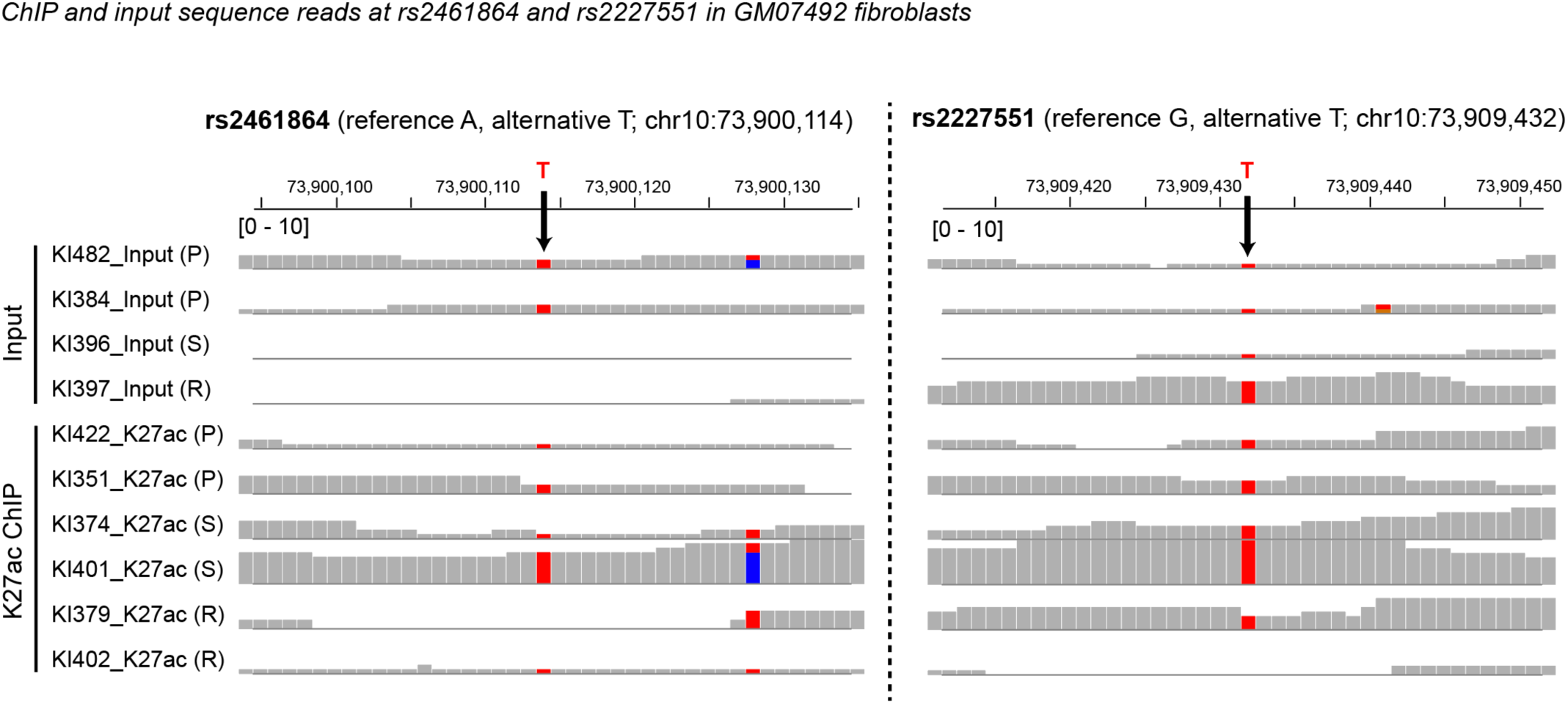
ChIP and input reads at rs2461864 and rs2227551 in GM07492 fibroblasts used in the serum starvation experiment. ChIP-seq and input sequencing reads from GM07492 are visualized using IGV. The T allele at both SNP sites is the IBD-risk allele. P, proliferation. S, starvation. R, recovery. Only the T (risk) allele was found at both locations, suggesting that GM07492 is homozygous for the risk allele.

**Supplementary Figure S5.**
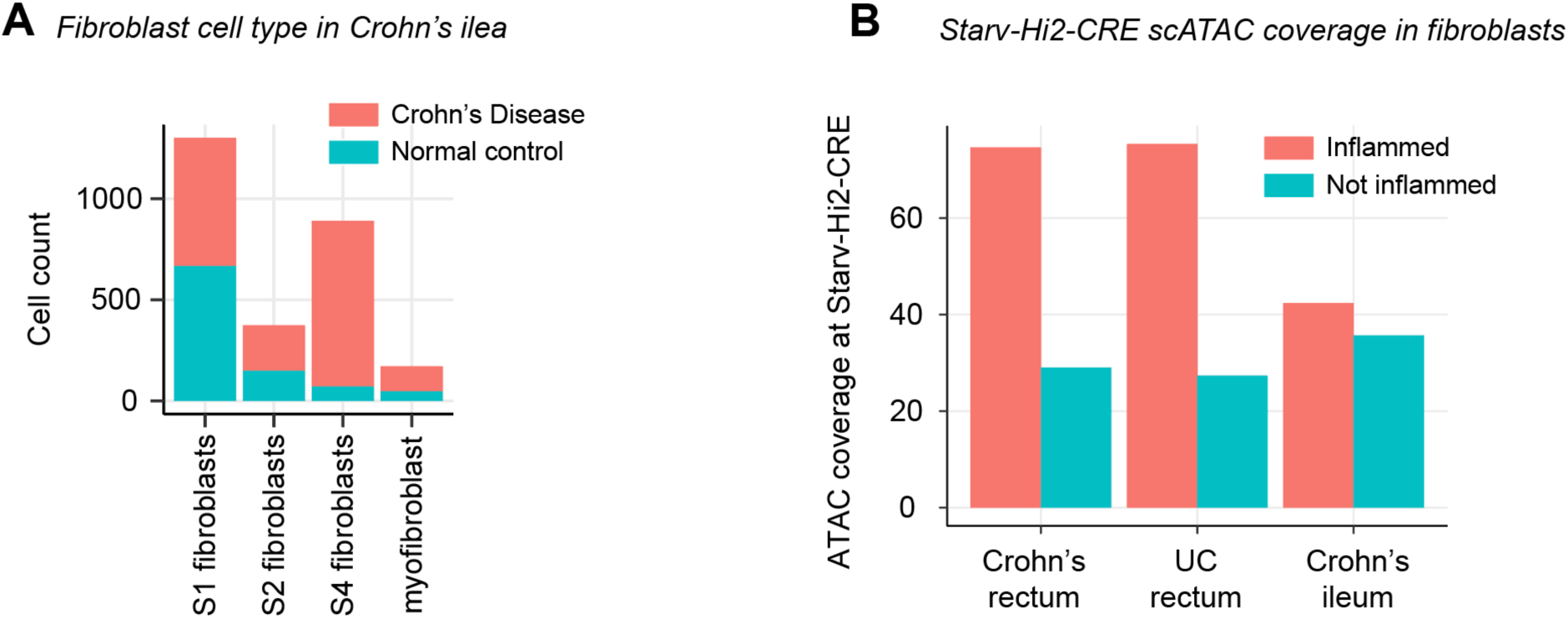
Additional analysis of scRNA-seq and scATAC-seq data for IBD intestines. **A.** Fibroblast cell count by fibroblast type in normal and Crohn’s disease ilea. Data are from Elmentaite et al. 2020. **B.** Quantification of scATAC-seq coverage at Starv-Hi2 CRE. Data are from Wayman et al. 2024.

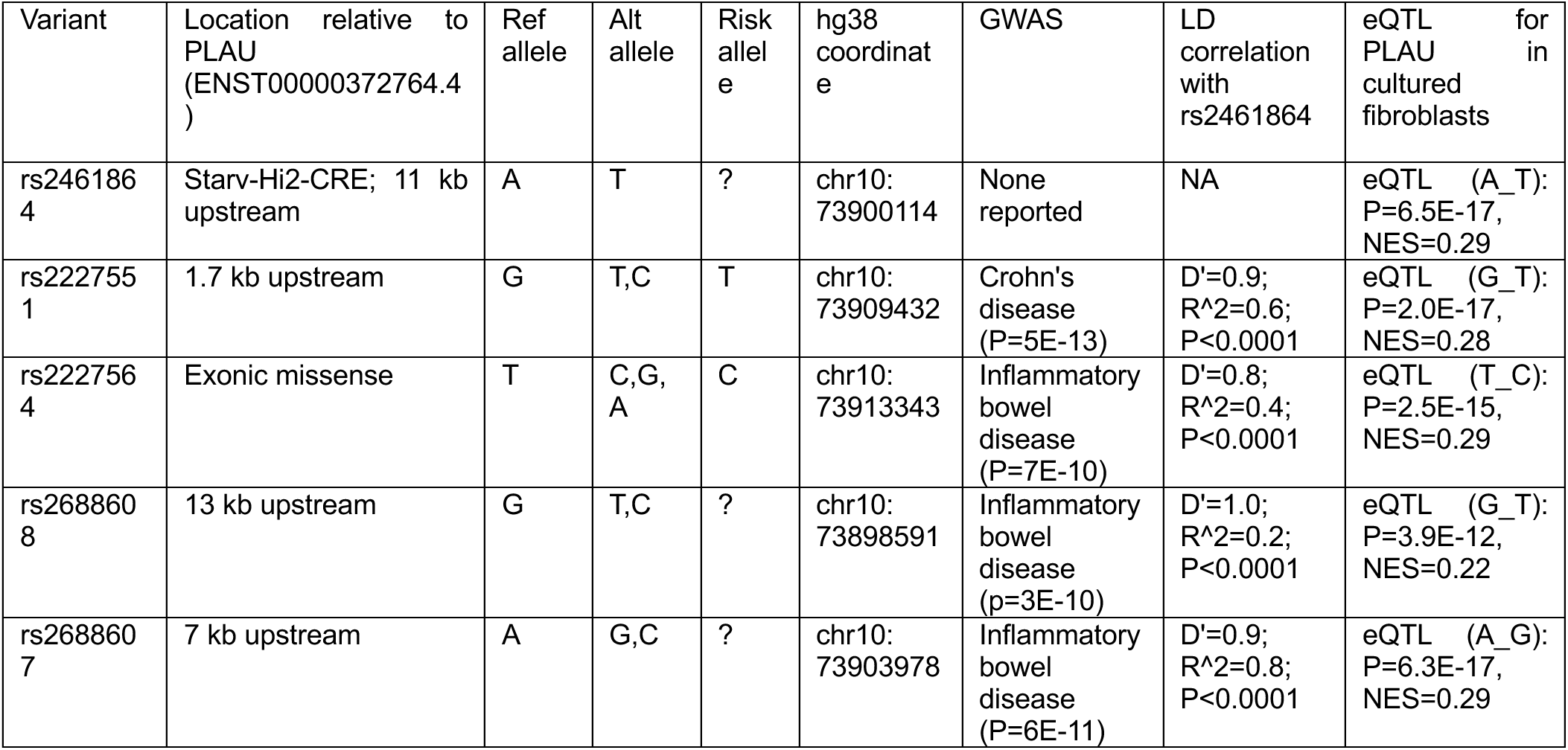

